# Functional Spatial Mapping of the Tumor Immune Microenvironment In Advanced Melanoma Patients

**DOI:** 10.1101/2025.05.29.656855

**Authors:** Emma Wagner, Samuel Legg, Christopher J Applebee, Amanda R Kirane, Julian Padget, Banafshé Larijani

## Abstract

**Introduction:** Current spatial proteomic approaches quantify immune checkpoint expression but do not directly measure functional receptor/ligand (PD-1/PD-L1) interactions within the tumor immune microenvironment (TiME). Therapeutic antibodies disrupt receptor-ligand interactions and do not target protein abundance. Methods that resolve functional checkpoint interactions provide biologically distinct insight beyond expression-based assays

**Methods:** We combined computation and quantitative spatial imaging, FuncO:TiME, [Functional Oncology Mapping (FuncOmap)], to map PD-1/PD-L1 interaction states to spatially defined regions of the TiME, in clinically annotated melanoma specimens, collected before and after neoadjuvant immune checkpoint blockade (ICB),

**Results:** FuncOmap spatially quantified millions of per-pixel PD-1/PD-L1 interactions demonstrated spatial heterogeneity in checkpoint interaction, not reflected by PD-1 expression levels alone. Post-treatment tissues exhibited increased PD-1/PD-L1 interaction states despite no corresponding increase in expression, indicating persistent or augmented functional checkpoint interaction despite therapy. Integration with spatial immune profiling further demonstrated that checkpoint interaction intensity can be contextualized within distinct immune cell populations.

**Conclusion:** We have established the feasibility of spatially resolved functional checkpoint mapping in human melanoma tissues. We demonstrate that receptor-ligand interactions diverge from protein expression patterns. By enabling direct interrogation of functional checkpoint interaction dynamics within intact tissue architecture, FuncO:TiME advances a functional paradigm for studying immune regulation in cancer.

## Introduction

Immune checkpoint blockade (ICB) has transformed the treatment landscape for advanced melanoma, yet clinical response remains heterogeneous, and robust biomarkers to guide therapeutic decision-making remain limited. Current tissue-based approaches, including digital spatial proteomics and transcriptomics, primarily quantify protein and RNA expression within the tumor immune microenvironment (TiME)^1–3^. While these methods have provided important insight into immune composition and checkpoint protein abundance, expression levels alone do not directly measure functional receptor-ligand interactions (e.g. downstream signaling impacted by molecular and conformational changes and post-translational modifications). Therapeutic antibodies target protein interactions rather than protein abundance, therefore inferring functional checkpoint interactions from expression-based metrics may obscure critical aspects of immune regulation, particularly in the context of ICB therapy^4,5^.

Time-resolved Förster resonance energy transfer (FRET) captured using fluorescence lifetime imaging microscopy (FLIM) quantifies intra- and intercellular receptor-ligand interactions within 1-10nm and has previously demonstrated that functional immune checkpoint interaction states can diverge from protein expression levels in multiple solid tumor types^6–11^.

The spatial resolution of intr**a**- and **i**ntercellular time-resolved FRET (**ai**FRET) has recently been further advanced through the development of FRET-based functional oncology mapping (FuncOmap), which directly quantifies and spatially localizes protein interaction states in fixed human tissues at per pixel resolution^6,10,12–15^. These aiFRET and FuncOmap studies underscore the conceptual distinction between checkpoint expression and checkpoint interaction and how the latter may associate with clinical outcomes more closely than abundance metrics alone^7,10–16^. Supplementary Table 1 provides a comparative overview of the advantages and limitations of aiFRET and FuncOmap relative to other antibody-based assays commonly used to quantify immune checkpoint interactions, including proximity ligation assay (PLA) and immunohistochemistry (IHC).

Although FuncOmap precisely quantifies receptor-ligand interaction states, it does not contextualize these interactions within defined TiME cell populations. Conversely, digital spatial profiling (DSP) platforms provide multiplexed, spatially resolved protein expression data that enable characterization of tumor and immune cell types expressing PD-1 and PD-L1, but lack the resolution required to determine which TiME cell populations contribute to functional interactions^1–3^.

To address these caveats, we developed FuncO:TiME; an imaging and computation framework that maps multiple frequency domain FLIM (mfFLIM)-based FuncOmaps and DSP to Hematoxylin & Eosin (H&E)-stained tissue sections to enable spatial mapping of PD-1/PD-L1 interaction states within clinically annotated melanoma tissues. This integration correlates per-pixel quantification of functional interactions with spatial immune cell characterization, permitting contextualization of checkpoint interaction intensity within distinct immune cell populations of the TiME.

In this methodological and feasibility study, we apply FuncO:TiME to melanoma specimens collected before and after neoadjuvant ICB. We demonstrate that PD-1/PD-L1 interaction states exhibit marked spatial heterogeneity that is not reflected by PD-1 or PD-L1 expression alone and illustrate post-treatment divergence in interaction patterns across clinically annotated response groups. These findings establish a framework for spatially resolved functional checkpoint mapping in human tumors and provide a foundation for future studies investigating checkpoint interaction dynamics in larger patient cohorts.

## Methods and Materials

### Antibodies and Reagents

See Supplementary Methods and Materials

#### Patient Samples

Formalin-fixed paraffin-embedded (FFPE) slides collected from patients with advanced melanoma were used in this study. Samples were collected prior to standard therapies and at the time of interval treatment assessment. All samples and clinical data were collected with the approval of Stanford University’s Institutional Review Board (IRB). Because this study was designed as a methodological feasibility investigation, formal power calculations were not performed. Patient response outcomes were blinded and only revealed when analyses were finalized. The clinicopathologic features of patients used in this study are summarized in Supplementary Table 2.

#### FuncOmap two-site assay for PD-1-PDL1 functional interactions

This follows already published methodology. Details in Supplementary Methods and Materials

#### Slide Preparation and COMET seqIF

Briefly, FFPE slides were de-waxed and antigen retrieval was performed by placing the slides in Dewax and HIER Buffer H (Epredia, catalogue number: TA-999-DHBH) inside the PT Module (Epredia, catalogue number: A80400011) for 20 mins at 95°C. Processed slides were immediately placed into Multi-staining Buffer (Lunaphore Technologies SA, catalogue number: BU06) until use. Slides were transferred to the COMET instrument where a microfluidic COMET Chip (Lunaphore Technologies SA, catalogue number: MK03) was sealed on top of the processed FFPE tissue section by the COMET instrument to form a closed reaction chamber. FFPE tissue sections were stained by reagents delivered via the microfluidic channels according to the pre-loaded seqIF protocols. Automated seqIF protocols were generated using the COMET Control software and consisted of iterative autofluorescence, staining, imaging and elution cycles^17^. Further details in Supplementary Methods and Materials.

#### COMET seqIF image analysis

Cell detection results across the tumor and FuncOmap regions (Ef 1 to 6) in Figure 4, were exported from QuPath as a csv file containing cell metadata, spatial coordinates and fluorescence intensity measurements for each antibody marker of the COMET seqIF panel. The csv file was uploaded to R (v4.5.2) using packages ‘read.csv’ and ‘data.table’ where mean fluorescence intensities for relevant markers were extracted for each segmented cell, and positivity thresholds were defined using 90^th^ percentile of fluorescence intensity for each marker within the dataset, a commonly used threshold for identifying the highest-expressing cell populations in spatial proteomic datasets. Spatial coordinates (centroid X/Y µm) were used to map each annotated cell to their location within the tumor region and were visualized using ‘dplyr’ and ‘ggplot2’

#### FuncO:TiME Analysis

This refers to spatial quantification of cells present within FuncOmap regions Ef 1 to 6. Ef region-specific spatial analyses were performed by sub-setting cell detection results to cells previously annotated as belonging to regions Ef 1 to 6 in QuPath. For each Ef region, cell co-ordinates (centroid X/Y µm) were plotted as two-dimensional scatter plots in R (ggplot2) where each dot corresponded to a single cell and was colored according to expression of PD-1/PD-L1 (i.e. PD status), immune cell lineage or TAM phenotype. Individual Ef regions were also displayed as their own scatter plot to help visualize region-specific distribution of cell PD status and immune lineage. For immune cell lineage plots across Ef 1 to 6, cells were initially designated as immune or melanoma cells based on 90^th^ percentile thresholds for CD45 (immune) or HMB45 +/-SOX10 (melanoma) markers respectively. Immune cells were subsequently classified into lineages based on passing 90^th^ percentile fluorescence intensity thresholds for the following marker combinations: tumor-associated macrophages (TAMs: CD68^+^), regulatory T cells (Tregs: CD3^+^CD4^+^FOXP3^+^), CD8 T cells (CD3^+^CD8^+^), CD4 T cell (CD3^+^ CD4^+^), CD3 T cell (CD3^+^), natural killer cells (NK: CD56^+^), NKT (CD3^+^CD56^+^) B cells (CD20^+^), dendritic cell (DC: CD11c^+^), monocyte (CD14^+^), neutrophils (CD16^+^CD15^−^), myeloid-derived suppressor cells (MDSC: CD16^-^CD15^+^), myeloid (CD11b^+^). For TAM-specific lineage plots, cells previously labelled as TAMs were sub-classified into M1-like, M2-like and M0-like TAMs based on the following marker combinations: CD68^+^CD80^+^ and/or CD86^+^ (M1-like), CD68^+^CD163^+^ and/or CD206^+^ (M2-like) and CD68^+^ only (M0-like). The proportion of immune cell lineages and TAM-phenotypes present in each Ef region were again visualized as bar plots using ggplot2.

#### Computational Framework workflow

FLIM-derived FRET efficiency (Ef) maps were generated using the FuncOmap software pipeline^10^, in which each pixel represents the calculated Ef value corresponding to PD-1/PD-L1 interaction. Ef maps were computationally aligned to clinically annotated H&E sections to enable spatial comparison between functional checkpoint interactions and histologic features. Alignment was performed using a five-step computational procedure. First, sampling paths connecting FLIM acquisition regions were reconstructed from microscope stage position metadata. Second, Otsu thresholding was applied to the H&E image to generate a tissue mask. Third, global scaling, rotation and translation were applied to maximize overlap between FLIM sampling regions and the H&E tissue mask. Fourth, local optimization was performed around each sampling region individually within a 200-pixel window (corresponding to approximately 10% of the H&E image dimensions). Finally, regions were identified where H&E tissue signal spatially coincided with acceptor fluorescence intensity measured by FuncOmap. Consequently, the user is presented with a map of points overlaid on the H&E image. Clicking on a point opens a heatmap of the variations in Ef for that region. On this heatmap, using the system’s pseudo-colormap, high Ef areas show white, while lower-intensity regions transition through yellow and red to black^29,30^. In addition, for each 512×512 pixel coincidence FLIM image frame, 1,000 datapoints were randomly sampled from 2.6×10^6^ total datapoints to generate mini violin plots representing the local distribution of Ef values. Sampling stability was evaluated by generating three independent random samples per region and confirming that their distributions were comparable. The resulting spatial map enables visualization of region-specific Ef distributions overlaid to histologic architecture. The code supporting image registration and interaction mapping will be made available upon reasonable academic request through the University of Bath.

### Statistical analyses

Violin plots intrinsically demonstrate the distribution of numerical data and are particularly effective for comparing multiple groups side-by-side. Hence, violin plots were used to visualize the distribution of Ef values across tissue regions and samples.

#### Mini violin plots

1,000 datapoints were randomly sampled from the total 2.6×10^6^ datapoints generated from each 512×512 pixel coincidence FLIM image frame as described above. This sample was used to generate a mini violin plot which provides a local approximation of the true distribution of Ef. The representativeness of the random samples was verified by generating three sample sets and checking that the variance between them was low.

#### Global violin plots

The global violin plots were generated from all 512×512 coincident images, corresponding to approximately 30×10^6^ datapoints per global violin plot. The upper quartile (Q3) of the global Ef distribution was used as a descriptive metric to represent regions with the highest PD-1/PD-L1 interaction states across tissues. Because Ef distributions were highly skewed and non-Gaussian, quartile-based summaries were used to visualize differences between samples.

## Results

This study was designed as a methodological feasibility analysis to develop and evaluate a computational framework that integrates histological annotation (H&E), functional interaction mapping (FuncOmap), and DSP to contextualize PD-1/PD-L1 interactions within distinct tumor regions and immune populations of the TiME.

### FuncOmap spatially quantifies PD-1/PD-L1 molecular engagement rather than inferring PD-1/PD-L1 proximity from protein abundance

The key differences in how traditional immunofluorescence (IF)-based assays and time resolved-FRET-based FuncOmap analysis quantify immune checkpoint interaction have previously been described in detail^10,11,16^ and are summarized in Figure 1. Briefly, IF is an example of a one-site assay where PD-1 and PD-L1 are independently labeled using conventional primary and secondary antibodies over consecutive cycles (Figure 1A). Conversely, FuncOmap is a spatial profile of a two-site assay where PD-1 and PD-L1 are labelled in the same cycle using conventional primary antibodies and species-specific F(ab’)2 fragments conjugated to either donor or acceptor chromophores (Figure 1D)^10,16^.

**Figure 1.**
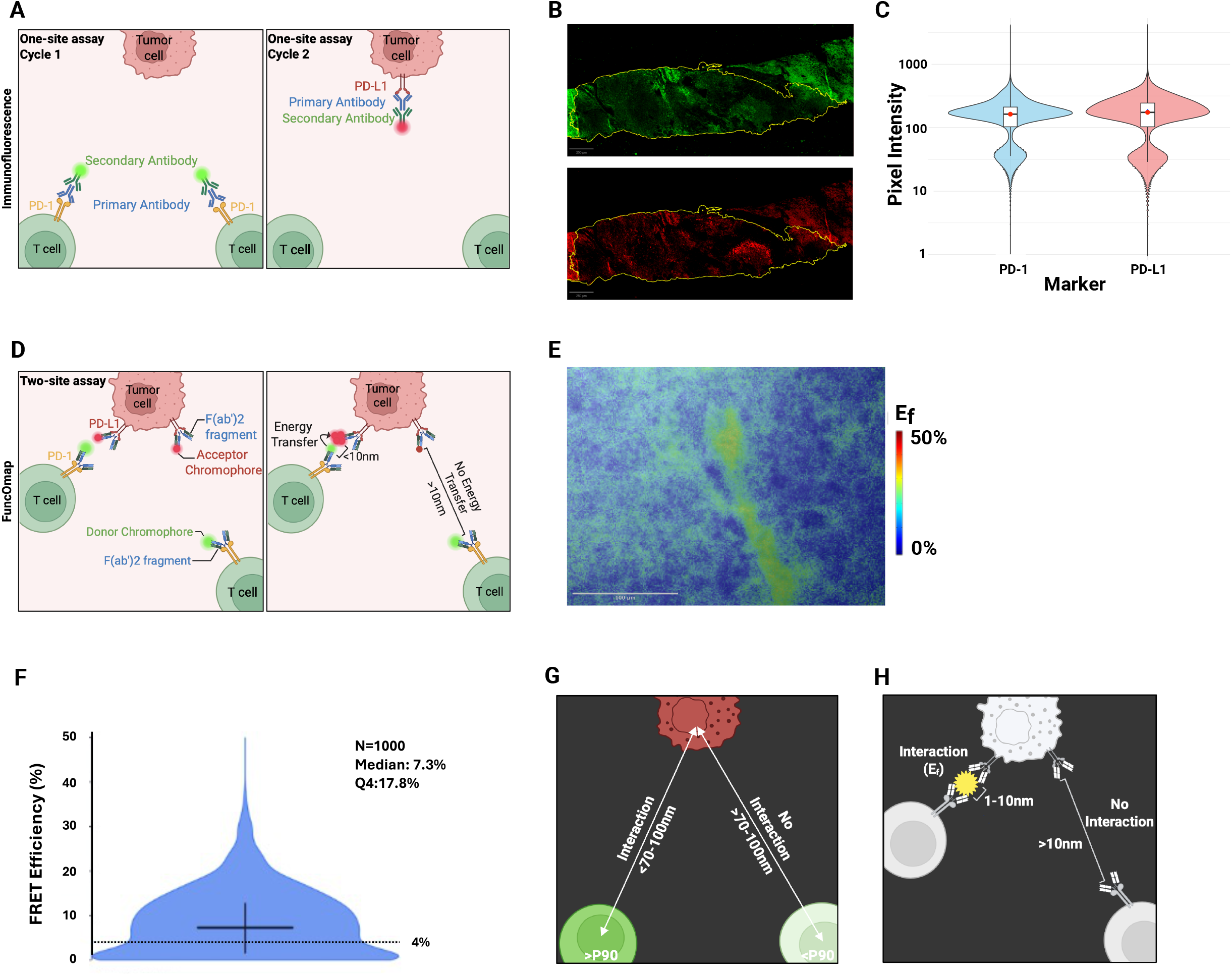
Spatial quantification of PD-1/PD-L1 interactions using one-site assays versus FuncOmap: A) Schematic depiction of a conventional one-site immunofluorescence assay; PD-1 and PD-L1 are labelled using a primary antibody followed by a fluorescent secondary antibody over independent cycles. Detection therefore reflects protein abundance at one antigen site per assay. B) COMET seqIF images showing spatial distribution of PD-1 (green) and PD-L1 (red) across a tumor-rich region of human melanoma tissue. C) Violin plots of the distribution of PD-1 and PD-L1 pixel intensities within the tumor region in B. D) Schematic depiction of two-site FuncOmap assay, where PD-1 and PD-L1 are labeled in the same cycle using donor- and acceptor-conjugated F(ab’2) fragments (green and red chromophores respectively). When chromophores are within 1-10nm, Förster resonance energy transfer (FRET) occurs. E) Representative FuncOmap heatmap from a single acquisition region (512×512 pixel frame) captured by multiple frequency domain fluorescence lifetime imaging microscopy (mfFLIM). Pixel color represents the FRET efficiency (Ef), reflecting the extent of PD-1/PD-L1 interaction. F) Violin plot showing the heterogeneous distribution of Ef values sampled from the region shown in panel E (n=1000 datapoints). The dotted line at 4% corresponds to the Förster radius (5.83nm) below which interaction is not detected. G) Conceptual illustration of interaction inference using expression-based profiling. PD-1^+^/PD-L1^+^ cells are annotated based on fluorescence intensity thresholds (90^th^ percentile, P90) and potential interactions are inferred from co-localization of PD-1^+^/PD-L1^+^ cells within a defined radius (70-100nm). H) Conceptual illustration of FuncOmap quantification of PD-1/PD-L1 interactions. PD-1/PD-L1 interactions are automatically quantified based on FRET between donor (PD-1) and acceptor (PD-L1) chromophores within 1-10nm of each other.

An example of COMET seqIF DSP of PD-1 and PD-L1 expression in stage III metastatic melanoma tissue is shown in Figure 1B. Regions of higher and lower PD-1 and PD-L1 expression are visually apparent and cells are often classified as PD-1^+^ or PD-L1^+^ using fluorescence intensity thresholds; with interactions inferred from their spatial proximity^22,23^. However, this binary classification does not capture the continuous variation in expression levels that influence receptor-ligand engagement. Consistent with this, the violin plot in Figure 1C demonstrates a heterogenous distribution of PD-1 and PD-L1 pixel intensities within an annotated tissue region.

FuncOmap enables spatial, per-pixel quantification of PD-1/PD-L1 interactions across tissue samples. These interactions are measured as FRET efficiency (Ef), where higher Ef values indicate close proximity and interaction between PD-1 and PD-L1 (Figure 1G)^10^. A representative heatmap illustrates the spatial heterogeneity of Ef values within a single FuncOmap region (Figure 1E), which is further captured by the corresponding violin plot (Figure 1F). Notably, Ef values below the Förster radius (R_0_) threshold (~4&) indicate a lack of interaction^10,16,24^.

### COMET seq IF and FuncOmap (FuncO:TiME) analyses are coordinated using clinically annotated H&E staining

The overall workflow from patient sample collection to FuncO:TiME analysis is summarized in Figures 2 and 3. Lymph node tissue samples from stage III metastatic melanoma patients were collected at baseline and after two cycles of neoadjuvant ICB treatment. H&E-stained sections were clinically annotated to define tumor and non-tumor regions, enabling coordinated spatial analysis across FuncOmap, COMET seqIF and downstream FuncO:TiME profiling.

**Figure 2.**
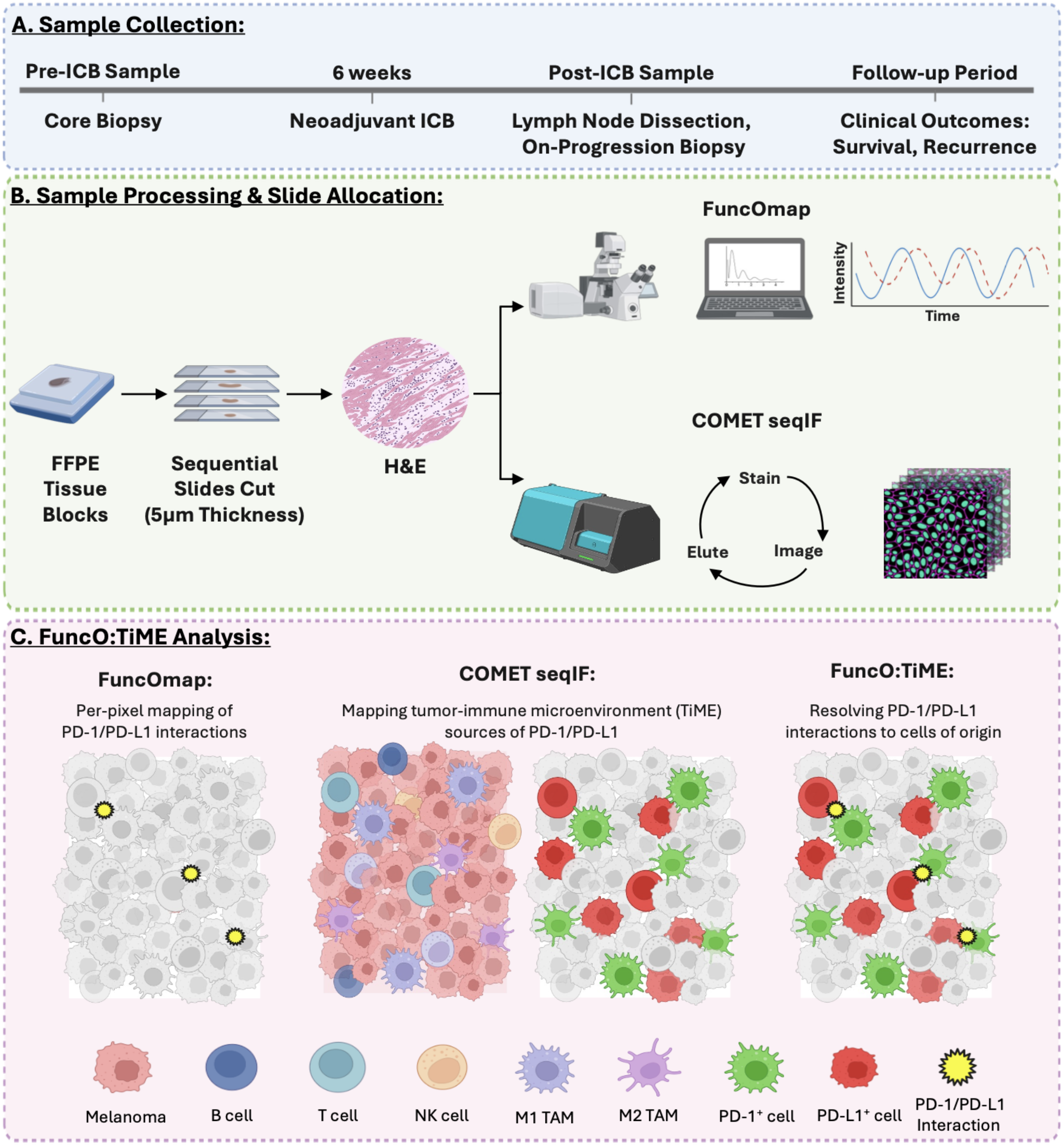
FuncO:TiME workflow integrating clinical sampling, spatial interaction mapping and immune cell profiling: A) Clinical sampling timeline. Tumor samples are obtained from melanoma patients prior to neoadjuvant immune checkpoint blockade (ICB) and again following standard therapy (~2 cycles) at time of lymph node dissection or on-progression biopsy. Clinical outcomes including recurrence and survival are assessed during follow-up. B) Sample processing and slide allocation. Formalin-fixed paraffin-embedded (FFPE) tissue blocks are sectioned into sequential slides (5µm thickness). Hematoxylin & Eosin (H&E)-staining is used for histologic annotation of tumor and non-tumor regions. Parallel sections are processed for FuncOmap acquisition and COMET seqIF to profile spatial immune cell populations through iterative staining, imaging, and antibody elution cycles. C) FuncO:TiME computational integration. FuncOmap generates per-pixel maps of PD-1/PD-L1 interactions across tissue sections, while COMET seqIF identifies tumor and immune cells within the tumor-immune microenvironment (TiME). The FuncO:TiME framework integrates these datasets to resolve functional PD-1/PD-L1 interactions to their cellular sources within the TiME.

**Figure 3.**
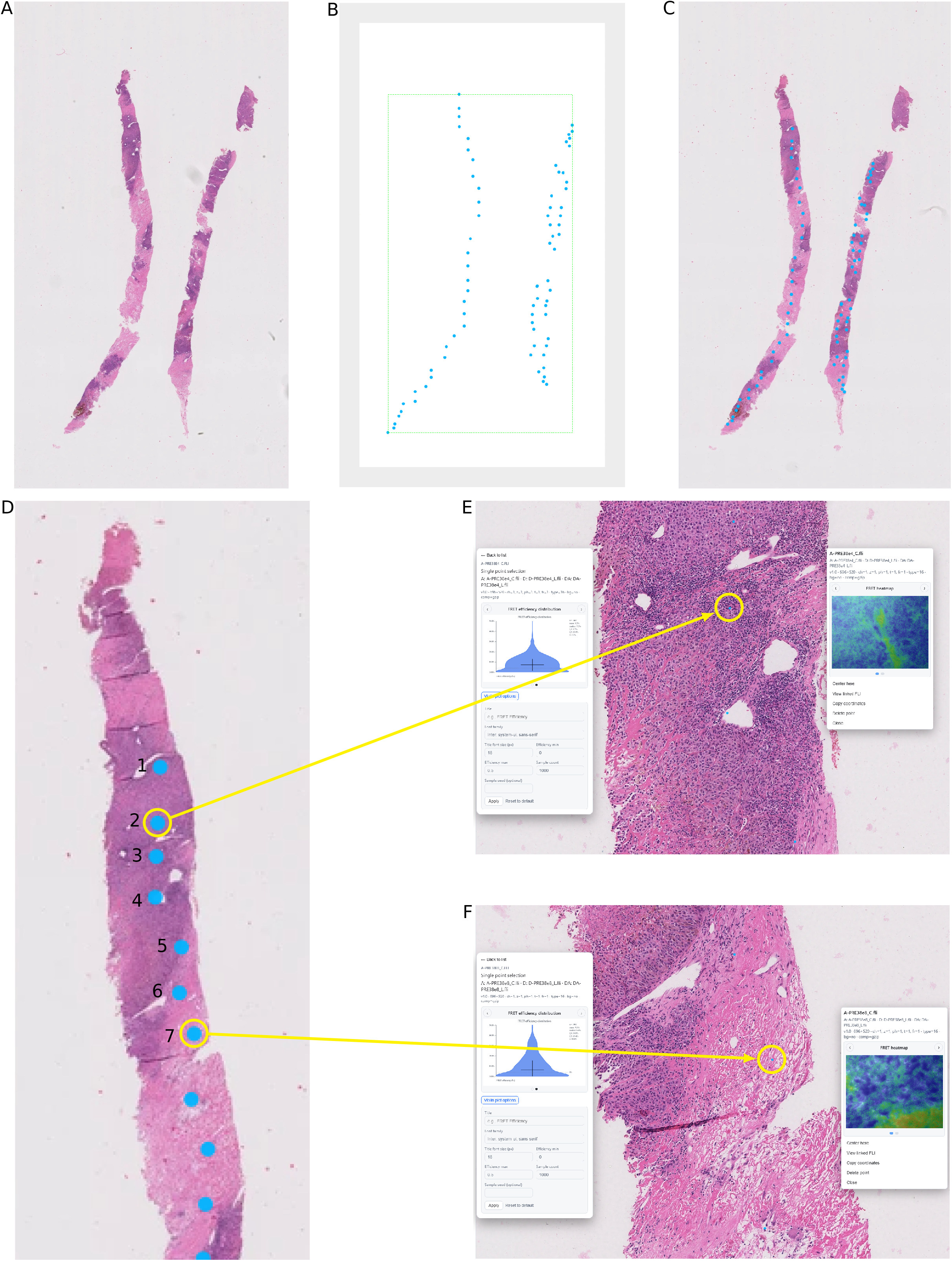
Spatial mapping of FuncOmap interaction measurements to histologic tissue architecture. This figure illustrates the computational workflow used to spatially register FuncOmap interaction measurements to histologically annotated tissue sections. Representative outputs (E-F) show regional analysis results for two example areas exhibiting high and low PD-1/PD-L1 interaction states. A) Whole slide image of an H&E-stained melanoma lymph node section used for histologic annotation. B) Spatial distribution of FuncOmap acquisition locations where mfFLIM quantifies PD-1/PD-L1 interaction efficiency (Ef). C) Overlay of FuncOmap acquisition coordinates onto the H&E tissue section, enabling spatial registration between interaction measurements and tissue morphology. D) Example mapping of individual FuncOmap acquisition regions along the tissue section. Numbered regions (1-7) represent selected areas used for regional analysis of PD-1/PD-L1 interaction efficiency (Ef). Individual points can be interrogated to visualize the regional Ef heatmap and distribution associated with that location. Ef values are visualized using a pseudo-color heatmap ranging from 0 to 50%, where higher interaction values appear white, and progressively lower values transition through yellow and red to black. E) Representative high-interaction region (tumor region 2, panel D) violin plot showing the distribution of pixel-level measurements within the FuncOmap acquisition area. F) Representative low-interaction region (non-tumor region 7) illustrating spatial heterogeneity in checkpoint interaction across the TiME and normal tissue interface.

**Figure 4.**
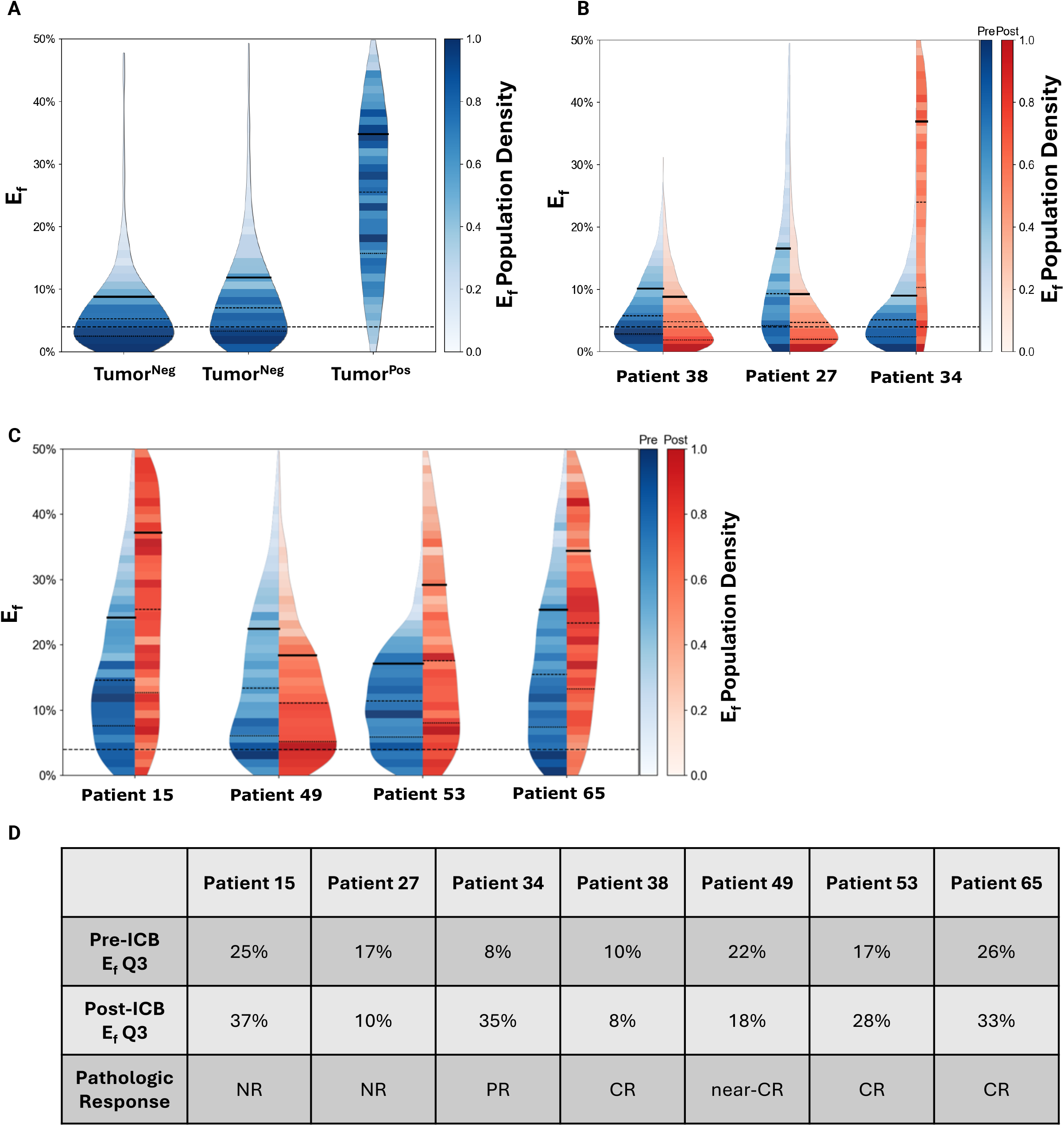
Global PD-1/PD-L1 interaction distributions across lymph node specimens before and after neoadjuvant immune checkpoint blockade (ICB). A) Baseline PD-1/PD-L1 interaction efficiency (Ef) distributions across control lymph nodes. The first tumor-negative (tumor^neg^) sample represents a lymph node from a stage IB melanoma patient without nodal metastasis. The second tumor^neg^ and tumor-positive (tumor^pos^) samples derive from a separate patient with one negative and one metastatic sentinel lymph node, enabling within-patient comparison of interaction states between tumor-free and tumor-involved tissues. Violin plots represent the distribution of pixel-level Ef measurements across tissue sections. B) Representative examples of global Ef distributions (30×10^6^ PD-1/PD-L1 Ef values) pre-(blue) and post-neoadjuvant ICB (red) in three melanoma patients with divergent pathologic responses (listed in panel D). C) Additional patient samples showing pre- and post-ICB Ef distributions across metastatic melanoma lymph node specimens. Each violin plot represents the distribution of pixel-level Ef across the analyzed tissue section D) Summary of Ef distribution changes across patients. The third quartile of the Ef distribution (Ef Q3) is shown for each patient pre- and post-ICB. Pathologic response classification is reported according to International Neoadjuvant Melanoma Consortium (INMC) criteria. (NR, non-response; PR, partial response; CR, complete response; near-CR; near-complete response). Within each violin plot, bold black lines indicate the Ef Q3, used as representative summary of global PD-1/PD-L1 interaction states within each tissue sample. Dotted horizontal lines at 4% Ef correspond to the Förster Radius (R_0_), representing the approximate threshold for detectable FRET, below which is considered non-interaction.

Mapping of FuncOmap regions to histologic features is illustrated in Figure 3. FuncOmap-derived Ef data were computationally aligned to annotated H&E sections, allowing spatial localization of PD-1/PD-L1 interactions within tissue architecture. This approach enabled region-specific integration of functional interaction data with histologic context.

### FuncOmap highlights differences in the distribution of PD-1/PD-L1 interactions between tumor and non-tumor regions and pre-versus post-ICI treatment

This study represents the first application of the current, functionally advanced version of FuncOmap to whole lymph node tumor samples from stage III metastatic melanoma patients pre- and post-ICB treatment.

In baseline lymph node samples from untreated stage IB melanoma patients undergoing routine sentinel lymph node biopsy (n=2), PD-1/PD-L1 interaction levels varied across tissue regions, with higher Ef values observed in tumor-positive (tumor^pos^) compared to tumor-negative (tumor^neg^) (Q3 Ef 35% versus 9-11%; Figure 4A).

Analysis of pre- and post-ICB lymph node tissues from stage III melanoma patients (n=7) similarly revealed marked heterogeneity in PD-1/PD-L1 interactions across samples (Figures 4B,C). However, no consistent differences in Q3 Ef values were observed across pathological response groups (CR, near-CR, PR, NR; Figure 4D).

### FuncOmap confirms PD-1/PD-L1 interactive states cannot be inferred from PD-1/PD-L1 relative abundance alone

Following the observation of higher PD-1/PD-L1 interactions in tumor regions, pre-ICB lymph node tissue from a complete responder (patient 38) was analyzed to assess spatial differences between tumor and non-tumor regions of a stage IIII melanoma sample (Figures 5A, B). Consistent with stage IB samples, PD-1/PD-L1 interactions were higher in tumor compared to non-tumor regions (Q3 Ef 15% versus 11%; Figure 5C).

**Figure 5.**
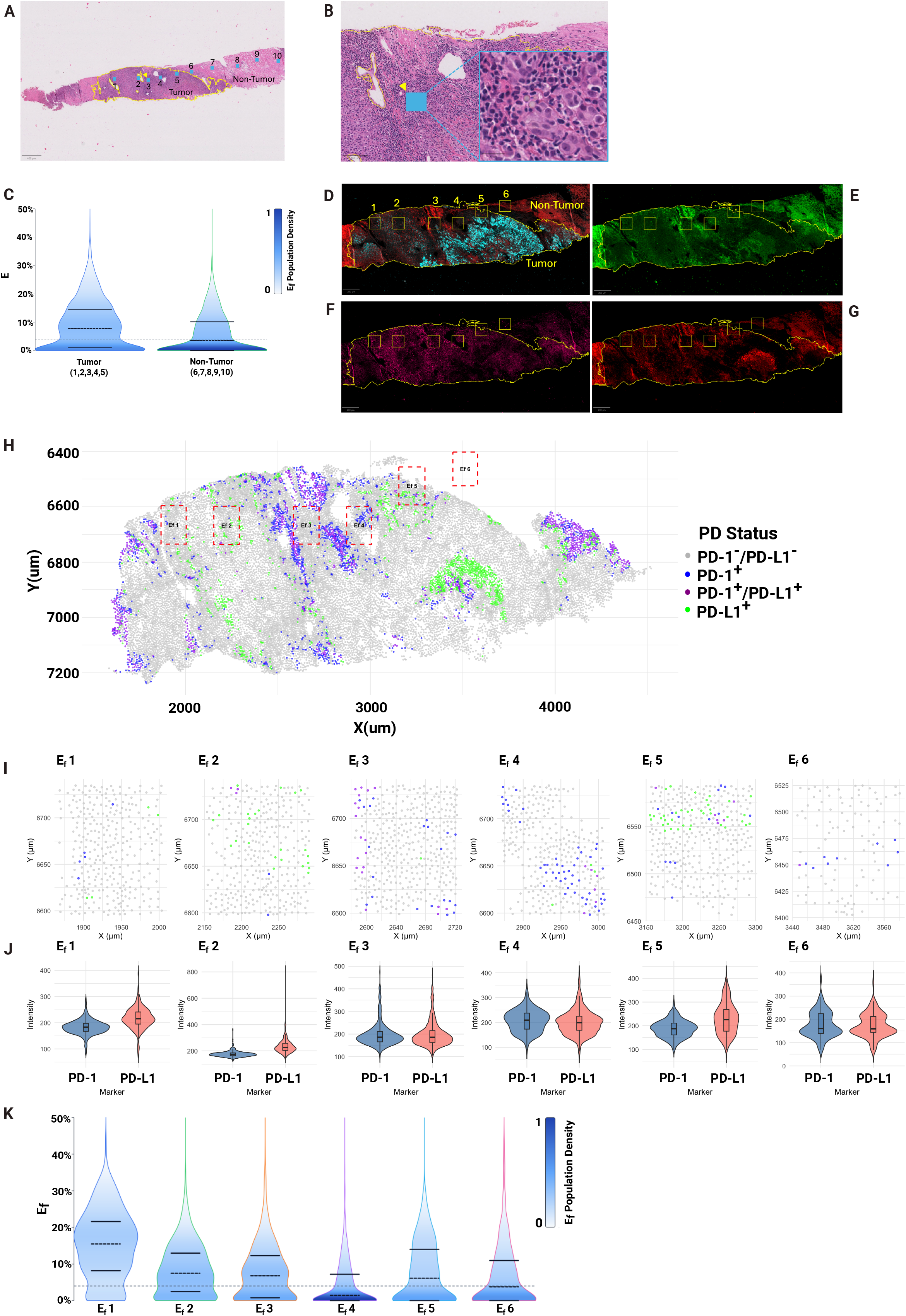
Integration of FuncOmap interaction mapping with spatial immune profiling using FuncO:TiME framework. A) Whole section H&E image of a representative melanoma lymph node metastasis (patient 38, pre-ICB) showing FuncOmap acquisition regions (blue boxes 1-10, identified using the workflow described in Figures 2-3) mapped across tumor (yellow boundary) and non-tumor regions. Regions 1-5 correspond to tumor and regions 6-10 correspond to adjacent non-tumor tissue. The scale bar represents 400µm. B) Higher magnification view of FuncOmap region 3 (blue box, highlighted by yellow arrow) from panel A. The scale bars represent 100µm (larger image) and 20µm (zoom view), respectively. C) Comparison of PD-1/PD-L1 interaction efficiency (Ef) distributions between selected tumor (regions 2-5) and non-tumor (9,10) FuncOmap regions. Upper (Q3) and lower (Q1) quartiles of Ef values are highlighted by bold solid black lines, while the median is highlighted by bold dotted black line. The dotted line at 4% corresponds to the detectable threshold for interaction. D) Spatial orientation of FuncOmap regions 1-6 integrated to COMET seqIF images (melanoma illustrated by HMB45^+^ [blue] and SOX10^+^ [red] cells. E-G) Spatial immune profiling of the same section using COMET seqIF. Marker distributions are shown across the tissue section to characterize protein and cellular abundance within the TiME (PD-1^+^ [green], CD45^+^ [pink], and PD-L1^+^ [red]). The scale bars represent 200µm. H) Spatial distribution PD-1^+^ and PD-L1^+^ expression at the single-cell level across the tissue section. FuncOmap regions (Ef 1-6) highlighted in red boxes, PD-1^-^/PD-L1^-^ (grey), PD-1^+^ (blue), PD-1^+^/PD-L1^+^ (purple) or PD-L1^+^ (green). I) Spatial distribution of cells within selected FuncOmap regions (Ef 1-6) used for downstream analysis. J) Quantification of PD-1 and PD-L1 fluorescence intensity of cells from each FuncOmap region (Ef 1-6). Violin plots represent distributions of marker intensity, reflecting relative protein abundance. K) Regional distributions of PD-1/PD-L1 interaction efficiency (Ef) across FuncOmap regions (Ef 1-6).

To examine how these interaction patterns related to protein abundance and cellular context, COMET seqIF analysis was performed on the same tissue. Spatial mapping of PD-1 and PD-L1 expression revealed heterogeneous distributions of PD-1^+^ and PD-L1^+^ cells across FuncOmap regions (Figure 5H-J).

Notably, PD-1/PD-L1 interaction levels did not correspond to relative protein abundance. The highest interaction levels were observed in Ef 1 (Q3 Ef 22%) despite a lower proportion of PD-1^+^ and PD-L1^+^ cells, whereas Ef 4 showed low interaction levels (Q3 Ef 7%) despite relatively higher proportion of PD-1^+^ and PD-L1^+^ cells (Figure 5K). Other regions showed comparable interaction levels despite differing expression patterns. Together, these findings demonstrate that PD-1/PD-L1 co-expression does not reliably reflect functional interaction, as captured by FRET-based FuncOmap analysis.

### FuncO:TiME reveals baseline tumor region PD-1/PD-L1 interactive states are associated with increased tumor associated macrophage infiltration

Having established Ef region-specific analyses, we next applied FuncO:TiME to examine how TiME cell distributions varied across Ef regions. Immune composition differed across regions (Figure 6A), with tumor-associated macrophages (TAMs) showing the strongest concordance with PD-1/PD-L1 interaction patterns. Higher TAM abundance was observed in regions with elevated PD-1/PD-L1 interaction (Ef 1, 2 and 5), whereas TAMs were largely absent in Ef 4, which exhibited the lowest interaction levels.

**Figure 6.**
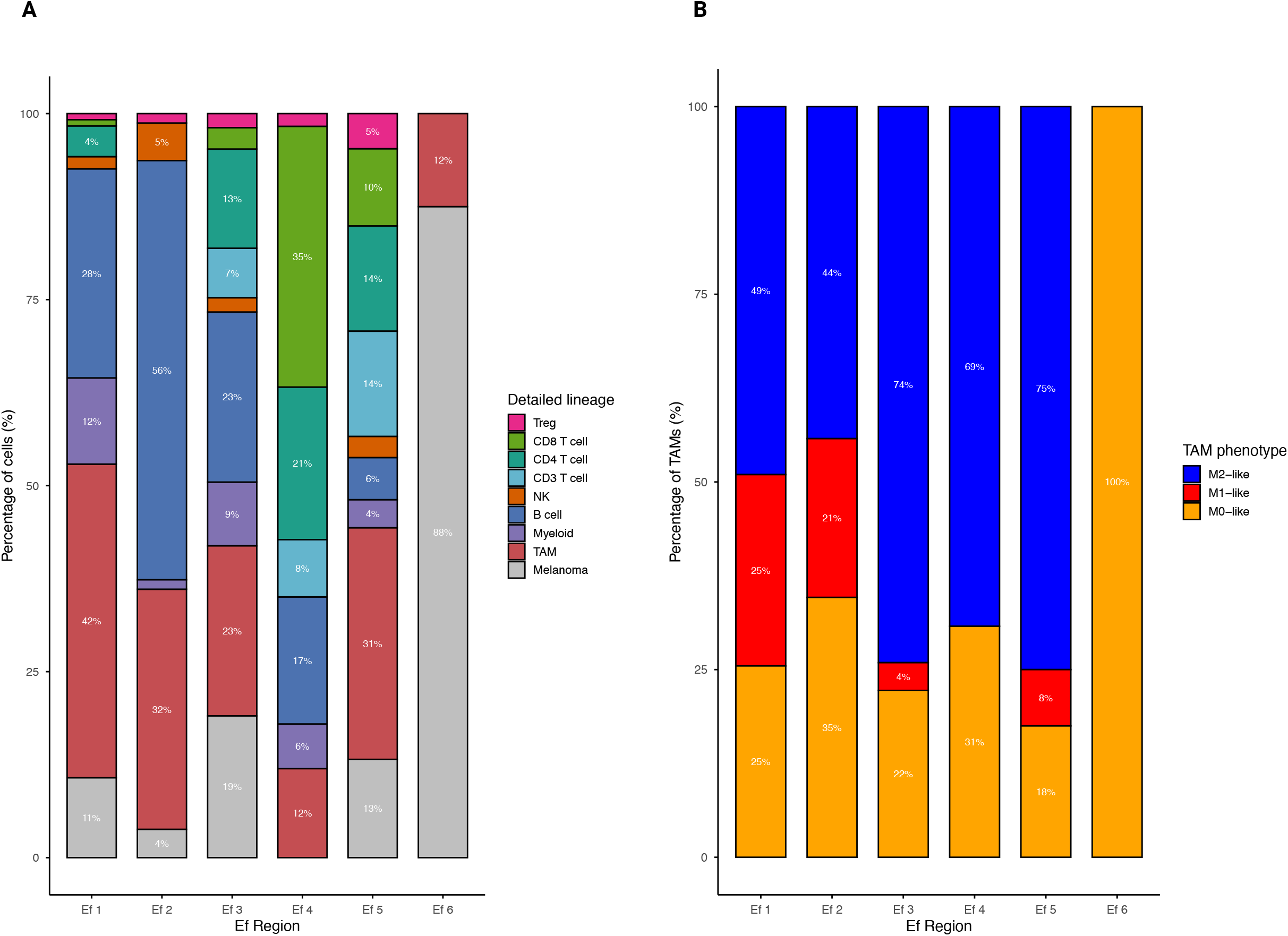
Immune cell lineage composition across FuncOmap interaction regions. A) Distribution of immune and tumor cell lineages across FuncOmap regions Ef 1-6 corresponding to spatial regions identified in Figure 5, representing areas with differing PD-1/PD-L1 interaction efficiencies. Stacked bars show the proportion of cells within each region classified as Regulatory T cells (Tregs), CD8 T cells, CD4 T cells, CD3 T cell, natural killer (NK) cells, B cells, Myeloid cells, Tumor Associated Macrophages (TAM), and Melanoma cells based on COMET seqIF lineage annotation. B) Distribution of TAM phenotypes across FuncOmap interaction regions (Ef1-6). Stacked bars show the proportion of M1-like, M2-like, and M0 macrophage populations within each region. TAM phenotypes were inferred from marker expression profiles obtained through COMET seqIF. These results suggest that variation in PD-1/PD-L1 interaction states may be associated with differences in TAM phenotype across tumor regions.

Further subclassification of TAM populations revealed distinct phenotypic patterns across Ef regions (Figure 6B). M1-like TAMs were enriched in regions with high PD-1/PD-L1 interactions (Ef 1 and 2), with moderate levels in Ef 3 and 5. In contrast, M2-like TAMs were most abundant in Ef 4, corresponding to low interaction levels, while Ef 6 was predominantly composed of M0-like TAMs. Together, these findings indicate that TAM abundance and phenotype track closely with PD-1/PD-L1 functional interactions.

## Discussion

The introduction of immune checkpoint blockade (ICB) to melanoma treatment has transformed outcomes for patients with advanced melanoma, yet durable responses remain limited and predictive biomarkers for patient selection remain imperfect^25^. Immunohistochemistry (IHC)-based clinical scoring of PD-L1 expression classifies tumors according to the proportion of PD-L1^+^ cells. However, such approaches infer checkpoint interaction indirectly from protein abundance and do not account for the spatial distribution, relative abundance or functional interaction state of PD-1 and PD-L1 within the TiME; all of which influence receptor-ligand engagment^26,27^. ICB acts by disrupting receptor-ligand interactions. Methods capable of directly resolving functional checkpoint interactions may provide biologically distinct information beyond expression-based assays.

In this study, we developed and applied a computational framework that integrates FuncOmap immune checkpoint interaction mapping with histologic annotation and digital spatial immune profiling to contextualize PD-1/PD-L1 interaction states within the TiME. Prior studies using intercellular FRET (iFRET) demonstrated that functional PD-1/PD-L1 interactions were detectable in tumors classified as PD-L1-negative by clinical assays and exhibit greater specificity than proximity-based methods such as PLA. However, these earlier iFRET approaches quantified interaction efficiency (Ef) without spatial localization within tissue architecture. **Func**tional **O**ncology **map**ping (FuncOmap) was therefore developed to extend iFRET analysis to spatially resolved tissue mapping, enabling regional characterization of functional checkpoint interactions across the TiME at the nanometer scale^10^.

Application of this approach to melanoma lymph node specimens demonstrated that PD-1/PD-L1 interactive states can diverge substantially from protein expression patterns. Regions with relatively low PD-1 or PD-L1 abundance sometimes exhibited higher interaction states than regions with higher protein expression, underscoring that checkpoint interaction cannot be reliably inferred from expression alone. These findings support the concept that functional receptor-ligand interactions represent a distinct biological parameter that may complement existing spatial proteomic approaches.

Using this framework, we compared global PD-1/PD-L1 interaction states across pre- and post-ICB lymph node tissues collected from stage III melanoma patients treated with neoadjuvant ICB and classified according to the international neoadjuvant melanoma consortium (INMC) response criteria. Across patients, global interaction distributions did not segregate consistently by pathological response criteria (CR, near-CR, PR, NR). In several cases, PD-1/PD-L1 interaction levels increased following therapy, including 2 of 3 patients with CR despite absence of residual viable tumor. Given that pathological response categories are defined by differing levels of residual viable tumor post-ICB treatment, these results reinforce the need for FuncOmap to differentiate PD-1/PD-L1 interactive states across tumor versus non-tumor tissue areas. This finding motivates the need to spatially resolve what remaining TiME features and cellular components contribute to these interaction patterns.

To first differentiate PD-1/PD-L1 interactive states between tumor and non-tumor tissue regions, FuncOmap was integrated with clinically annotated Hematoxylin & Eosin (H&E) sections. These annotations were used to spatially assign FuncOmap acquisition regions to tumor and non-tumor tissue areas, enabling direct comparison of interactive states across distinct compartments of the TiME. Application of this approach to baseline lymph node tissue demonstrated higher PD-1/PD-L1 interaction levels within tumor-positive regions compared to tumor-negative regions. Similar compartment-specific differences were also observed in metastatic melanoma specimens. This improved FuncOmap methodology was applied to pre-ICB tissue collected from patient 38 (CR). Once again, global PD-1/PD-L1 interactive states were higher in tumor regions compared to non-tumor regions.

Recent multimodal spatial analysis pipelines combining multiplex immunofluorescence, imaging mass cytometry, and histologic annotation have demonstrated the potential to integrate molecular and morphological features within tissue sections^28^. However, integrating imaging data across these modalities with differing optical properties and acquisition systems remains technically challenging. In contrast, the FuncOmap computational framework enables automated mapping of tens of millions (30×10^6^ per-pixel) nanoscale PD-1/PD-L1 interaction states across entire tissue sections and computational alignment with H&E images, enabling functional checkpoint interaction mapping within defined TiME compartments.

Although the H&E staining and FuncOmap analyses presented in this paper were performed on two sequential slides, both methods are non-destructive and enable spatial alignment of interaction measurements with histologic architecture. Future methodological refinements may further optimize multimodal integration with spatial immune profiling platforms such as COMET seqIF while minimizing the number of sequential sections required. Integration with multiplex immune profiling further demonstrated that PD-1/PD-L1 interaction states cannot be reliably inferred from relative protein abundance alone and provides orthogonal spatial immune profiling that independently contextualizes FuncOmap interaction measurements.

Fluorescence intensities of PD-1 and PD-L1 were quantified across the tumor region to approximate relative protein abundance. Cells with the highest surface density were annotated using 90^th^ percentile intensity thresholds. However, FuncOmap analysis revealed that regions with lower apparent PD-1/PD-L1 expression sometimes exhibited higher interaction states than regions with greater expression. These observations reinforce the distinction between protein expression and functional receptor-ligand interaction and highlight the importance of directly measuring checkpoint interaction states when interpreting spatial immune signaling within the TiME. Direct measurement of checkpoint interaction may therefore provide complementary biological information to existing expression-based assays when interpreting immune checkpoint activity within tumor tissues.

Integration with spatial immune profiling further suggested that variation in checkpoint interaction states may be associated with differences in immune cell composition across tumor regions. Tumor-associated macrophages (TAMs) distribution most closely paralleled regional variation in PD-1/PD-L1 interaction levels in the specimens analyzed. Although CD8^+^ T cells are commonly considered key mediators of response to ICB, our exploratory analysis identified relatively limited CD8^+^ T cell presence within several high-interaction regions. Instead, variation in M1-like and M2-like TAM populations appeared to correspond with differences in interaction states. These results were of particular interest given our prior findings that baseline post-treatment enrichment of M1-like and M2-like TAMs were also associated with pathological responses to neoadjuvant immunotherapy with oncolytic virus talimogene laherparepvec (TVEC), respectively, in stage II melanoma patients^11^. Altogether, these observations are hypothesis-generating and require validation in larger cohorts but suggest TAMs may contribute to checkpoint interaction dynamics within the melanoma TiME.

Several limitations should be acknowledged. This study was designed as a methodological feasibility analysis and was conducted in a relatively small cohort of clinically annotated melanoma specimens. Consequently, the present findings should not be interpreted as establishing predictive biomarkers for immunotherapy response. In addition, spatial registration between imaging modalities currently relies on sequential tissue sections and histologic landmarks, which may introduce minor alignment variability. Ongoing methodological development aims to further refine image registration and expand application of the FuncO:TiME framework to larger patient cohorts.

## Conclusion

The FuncO:TiME framework provides a platform for spatially resolved mapping of functional checkpoint interactions within intact tumor tissues. By enabling direct quantification of PD-1/PD-L1 molecular interactions and contextualizing these interactions within the TiME, this approach establishes a foundation for future studies investigating the biological and clinical relevance of functional checkpoint interactions in cancers. Future work will focus on expanding this analysis to larger clinically annotated patient cohorts and further refining computational alignment between FuncOmaps and multimodal spatial profiling platforms to help inform the future development of functional biomarkers for patient selection.

## Acknowledgements

This manuscript is dedicated to the late Professor Laridjani, M. (1935-2025), whose tireless perseverance in the righteous path of scientific research became his way of being.

This work was generously supported by the John and Marva Warnock Endowed Scholar Fund, as well as by the Stanford Cancer Institute, an NCI- designated Comprehensive Cancer Centre and the Stanford Department of Surgery Equipment Grant

We would like to thank Hsiu-Ju Hsu (Sherry) for her help with patient sample preparations, classifications and optimizations of the Lunaphore-COMET.

## Notes

### Competing Interest Statement

The authors have declared no competing interest.

### Summary of Updates

Functional Oncology mapping (FuncOmap) and tumour immune microenvironment analysis of 7 more patients have been added. Control samples have been added. New version of FuncOmap used to determine more precise spatial analysis All Figures are new in the revised version

## References

1. Su DG, Schoenfeld DA, Ibrahim W, et al. Digital spatial proteomic profiling reveals immune checkpoints as biomarkers in lymphoid aggregates and tumor microenvironment of desmoplastic melanoma. J Immunother Cancer. 2024;12(3):e008646. doi:10.1136/jitc-2023-008646

2. Horowitch B, Lee DY, Ding M, et al. Subsets of IFN Signaling Predict Response to Immune Checkpoint Blockade in Patients with Melanoma. Clin Cancer Res Off J Am Assoc Cancer Res. 2023;29(15):2908–2918. doi:10.1158/1078-0432.CCR-23-0215

3. Aung TN, Warrell J, Martinez-Morilla S, et al. Spatially Informed Gene Signatures for Response to Immunotherapy in Melanoma. Clin Cancer Res Off J Am Assoc Cancer Res. 2024;30(16):3520–3532. doi:10.1158/1078-0432.CCR-23-3932

4. Martinez-Morilla S, Moutafi M, Fernandez AI, et al. Digital spatial profiling of melanoma shows CD95 expression in immune cells is associated with resistance to immunotherapy. Oncoimmunology. 2024;12(1):1. doi:10.1080/2162402X.2023.2260618

5. Zerdes I, Matikas A, Mezheyeuski A, et al. Machine learning-based spatial characterization of tumor-immune microenvironment in the EORTC 10994/BIG 1-00 early breast cancer trial. Npj Breast Cancer. 2025;11(1):23. doi:10.1038/s41523-025-00730-1

6. Sánchez-Magraner L, Miles J, Baker CL, et al. High PD-1/PD-L1 Checkpoint Interaction Infers Tumor Selection and Therapeutic Sensitivity to Anti-PD-1/PD-L1 Treatment. Cancer Res. 2020;80(19):4244–4257. doi:10.1158/0008-5472.CAN-20-1117

7. Larijani B, Miles J, Ward SG, Parker PJ. Quantification of biomarker functionality predicts patient outcomes. Br J Cancer. 2021;124(10):1618–1620. doi:10.1038/s41416-021-01291-3

8. Miles J, Ward SG, Larijani B. The fusion of quantitative molecular proteomics and immune-oncology: a step towards precision medicine in cancer therapeutics. FEBS Lett. 2022;596(21):2721–2735. doi:10.1002/1873-3468.14480

9. Miles J, Soubeyran I, Marliot F, et al. Determination of Interactive States of Immune Checkpoint Regulators in Lung Metastases after Radiofrequency Ablation. Cancers. 2022;14(23):5738. doi:10.3390/cancers14235738

10. Safrygina E, Applebee C, McIntyre A, Padget J, Larijani B. Spatial functional mapping of hypoxia inducible factor heterodimerisation and immune checkpoint regulators in clear cell renal cell carcinoma. BJC Rep. 2024;2(1):1. doi:10.1038/s44276-023-00033-7

11. Kirane AR, Lee D, Lowe M, et al. Toward Functional Biomarkers of Response to Neoadjuvant Oncolytic Virus in Stage II Melanoma: Immune-Förster Resonance Energy Transfer and the Dynamic Tumor Immune Microenvironment. JCO Oncol Adv. Published online January 2025. doi:10.1200/OA-24-00049

12. Kong A, Leboucher P, Leek R, et al. Prognostic value of an activation state marker for epidermal growth factor receptor in tissue microarrays of head and neck cancer. Cancer Res. 2006;66(5):2834–2843. doi:10.1158/0008-5472.CAN-05-2994

13. Kong A, Calleja V, Leboucher P, Harris A, Parker PJ, Larijani B. HER2 Oncogenic Function Escapes EGFR Tyrosine Kinase Inhibitors via Activation of Alternative HER Receptors in Breast Cancer Cells. PLOS ONE. 2008;3(8):e2881. doi:10.1371/journal.pone.0002881

14. Veeriah S, Leboucher P, de Naurois J, et al. High-throughput time-resolved FRET reveals Akt/PKB activation as a poor prognostic marker in breast cancer. Cancer Res. 2014;74(18):4983–4995. doi:10.1158/0008-5472.CAN-13-3382

15. Miles J, Applebee CJ, Leboucher P, et al. Time resolved amplified FRET identifies protein kinase B activation state as a marker for poor prognosis in clear cell renal cell carcinoma. BBA Clin. 2017;8:97–102. doi:10.1016/j.bbacli.2017.10.002

16. Wagner E, Larijani B, Kirane AR. Predictive Biomarkers for Immune Checkpoint Inhibitor Therapy in Advanced Melanomas. Surg Oncol Clin N Am. 2025;34(3):437–451. doi:10.1016/j.soc.2025.01.006

17. Navikas V, Kowal J, Rodriguez D, et al. Semi-automated approaches for interrogating spatial heterogeneity of tissue samples. Sci Rep. 2024;14(1):5025. doi:10.1038/s41598-024-55387-w

18. Bankhead P, Loughrey MB, Fernández JA, et al. QuPath: Open source software for digital pathology image analysis. Sci Rep. 2017;7(1):16878. doi:10.1038/s41598-017-17204-5

19. Schmidt U, Weigert M, Broaddus C, Myers G. Cell Detection with Star-Convex Polygons. In: Frangi AF, Schnabel JA, Davatzikos C, Alberola-López C, Fichtinger G, eds. Medical Image Computing and Computer Assisted Intervention – MICCAI 2018. Springer International Publishing; 2018:265–273. doi:10.1007/978-3-030-00934-2_30

20. Weigert M, Schmidt U, Haase R, Sugawara K, Myers G. Star-convex Polyhedra for 3D Object Detection and Segmentation in Microscopy. In: 2020 IEEE Winter Conference on Applications of Computer Vision (WACV). 2020:3655–3662. doi:10.1109/WACV45572.2020.9093435

21. Weigert M, Schmidt U. Nuclei instance segmentation and classification in histopathology images with StarDist. In: 2022 IEEE International Symposium on Biomedical Imaging Challenges (ISBIC). 2022:1–4. doi:10.1109/ISBIC56247.2022.9854534

22. Jin S, Guerrero-Juarez CF, Zhang L, et al. Inference and analysis of cell-cell communication using CellChat. Nat Commun. 2021;12(1):1088. doi:10.1038/s41467-021-21246-9

23. Hou R, Denisenko E, Ong HT, Ramilowski JA, Forrest ARR. Predicting cell-to-cell communication networks using NATMI. Nat Commun. 2020;11(1):5011. doi:10.1038/s41467-020-18873-z

24. Jares-Erijman EA, Jovin TM. FRET imaging. Nat Biotechnol. 2003;21(11):1387–1395. doi:10.1038/nbt896

25. Tong J, Kartolo A, Yeung C, Hopman W, Baetz T. Long-Term Toxicities of Immune Checkpoint Inhibitor (ICI) in Melanoma Patients. Curr Oncol. 2022;29(10):7953–7963. doi:10.3390/curroncol29100629

26. Paver EC, Cooper WA, Colebatch AJ, et al. Programmed death ligand-1 (PD-L1) as a predictive marker for immunotherapy in solid tumours: a guide to immunohistochemistry implementation and interpretation. Pathology (Phila). 2021;53(2):141–156. doi:10.1016/j.pathol.2020.10.007

27. Marletta S, Fusco N, Munari E, et al. Atlas of PD-L1 for Pathologists: Indications, Scores, Diagnostic Platforms and Reporting Systems. J Pers Med. 2022;12(7):1073. doi:10.3390/jpm12071073

28. Marlin MC, Stephens T, Wright C, Smith M, Wright K, Guthridge JM. A novel process for H&E, immunofluorescence, and imaging mass cytometry on a single slide with a concise analytics pipeline. Cytometry A. 2023;103(12):1010–1018. doi:10.1002/cyto.a.24789

29. Hunter JD. Matplotlib: A 2D Graphics Environment. Comput Sci Eng. 2007;9(3):90–95. doi:10.1109/MCSE.2007.55

30. Colormap reference — Matplotlib 3.10.8 documentation. Accessed March 13, 2026. https://matplotlib.org/stable/gallery/color/colormap_reference.html

