## Supplementary Methods for "Functional Spatial Mapping of the Tumor Immune Microenvironment In Advanced Melanoma Patients"

### Methods and Materials

#### Antibodies and Reagents

*FuncOmap:*

Monoclonal antibodies mouse anti-PD-1(AB_2619745), rabbit anti-PD-L1 (AB_2619745) were purchased from Abcam (catalogue numbers ab52587 and ab205921 respectively). AffiniPure F(ab')$_2$ fragment donkey anti-mouse IgG (H+L) and peroxidase AffiniPure F(ab')$_2$ fragment donkey anti-rabbit IgG (H+L) were purchased from Jackson Immuno Research (catalogue numbers 715-006-150 and 711-036-152 respectively). Pierce endogenous peroxidase suppressor, TSA SuperBoost kit and Prolong Glass antifade mount were purchased from Thermofisher Scientific (catalogue numbers 35000, B40925 and P36980 respectively). ATTO488 NHS ester, bovine serum albumin and rhodamine B, were purchased from Sigma Aldrich (catalogue numbers A2153-100G and 234141-10G respectively).

*COMET Sequential immunofluorescence (seqIF):*

| Marker | Host species | Company, Clone | Catalogue Number: | Dilution |
|---|---|---|---|---|
| CD45 | Mouse | Lunaphore, (PD7/26 + 2B11) | MR10090 | 1:100 |
| Cd11b | Rabbit | Novus, (polyclonal) | NB110-89474 | 1:100 |
| HMB45 | Mouse | Abcam, (HMB45 + M2-7C10 + M2-9E3 + T311) | Ab733 | 1:100 |
| CD11c | Rabbit | Lunaphore, (BLR138H) | MR10070 | 1:100 |
| CD56 | Mouse | Invitrogen (123c3) | MA1-06801 | 1:100 |
| CD68 | Mouse | Abcam, (KP-1) | Ab955 | 1:200 |
| CD14 | Rabbit | Proteintech, (polyclonal) | 17000-1-AP | 1:200 |
| CD15 | Rabbit | Abcam, (SP159) | Ab135377 | 1:100 |
| CD20 | Mouse | Abcam, (L26) | Ab9475 | 1:100 |
| CD8 | Mouse | Lunaphore (C8/144B) | MR10030 | 1:200 |
| CD3e | Rabbit | Abcam, (polyclonal) | ab5690 | 1:100 |
| CD4 | Rabbit | Lunaphore, (BL-155-1C11) | MR10020 | 1:100 |
| FoxP3 | Mouse | Lunaphore, (236A/E7) | MR10040 | 1:100 |
| CD80 | Mouse | Invitrogen, (2A2) | MA5-15512 | 1:50 |

| | | | | |
|---|---|---|---|---|
| CD86 | Rabbit | Abcam (EPR21962) | Ab239075 | 1:200 |
| CD163 | Rabbit | Abcam, (EPR14643-36) | Ab218294 | 1:200 |
| CD206 | Rabbit | Proteintech, (polyclonal) | 18704-1-AP | 1:200 |
| MHCII | Mouse | Abcam, (6C6) | ab55152 | 1:200 |
| HLA-DR | Mouse | Abcam, (TAL 1B5) | ab20181 | 1:200 |
| IBA-1 | Rabbit | Abcam, (EPR16588) | Ab178846 | 1:10,000 |
| AXL | Rabbit | Cell Signaling Technologies, (C89E7) | 8661S | 1:200 |
| PD-1 | Rabbit | Abcam, (NAT105) | Ab52587 | 1:100 |
| PD-L1 | Rabbit | Abcam, (28-8) | Ab205921 | 1:100 |
| CTLA4 | Rabbit | Invitrogen, (11H7L17) | 702534 | 1:50 |
| LAG3 | Rabbit | Cell Signaling Technologies, (D2G4O) | 15372S | 1:50 |
| Secondary antibodies: | | | | |
| Donkey anti-mouse IgG (H+L) Alexa Fluor 555 | | Invitrogen | A-31570 | 1:2000 |
| Goat anti-Rabit IgG (H+L) Alexa Fluor 647 | | Invitrogen | A-21244 | 1:4000 |
| Nuclear Staining: | | | | |
| Hoechst 33342 | | Invitrogen | H3570 | 1:1000 |

**FuncOmap two-site assay for PD-1-PDL1 functional interactions**

The FFPE slides underwent antigen retrieval process using the Envision Flex retrieval solution, pH 9. The Dako PT-Link system was utilized, where the slides were heated to 95°C for 20 minutes. Using a PAP pen, an aqueous-repelling border was outlined around each tissue fragment. Pierce endogenous peroxidase suppressor was then applied to each specimen, and the slides were left to incubate for 30 minutes at 21°C in a humidity-controlled environment. The samples underwent two washes with PBS before being incubated for an hour at room temperature with 3% BSA (10 mg/ml). For the donor-only slides, these were incubated with anti-PD-1 (at a dilution of 1:100).

The donor-acceptor slides were treated with the following primary antibodies: αPD-1 (1:100) and αPD-L1 (1:500). Primary antibodies were incubated overnight at 4°C. Samples were washed with 0.02% PBST. The samples were treated with secondary F(ab')2 fragments. F(ab')2- ATTO488 (at a dilution of 1:100) was introduced to the donor-only slides. As for the donor-acceptor slides, they received both F(ab')2-ATTO488 (1:100) and F(ab')2-horseradish peroxidase (HRP) (1:200). These samples were incubated for 2 hours in the dark, at room temperature in a humidified container. After the incubation period slides were washed with 0.02% PBST. The donor-only slides were mounted with 1 drop of Prolong Glass antifade mount. The donor-acceptor slides were subjected to tyramide signal amplification (TSA). The purpose of TSA was to amplify the acceptor labelling, thus increase the signal-to-noise ratio and enhancing resonance energy transfer. This procedure is described in detail in Veeriah et al., 2014[14] and Magraner Sanchez et al., 2020[6]. The antibodies were labelled with species-specific $F(ab')_2$ fragments which were conjugated to ATTO488 (donor chromophore, used to label the receptor primary antibody) or HRP (used to label the ligand primary antibody). TSA was used to conjugate the acceptor chromophore (Alexa594) to the HRP labelling the ligand. This method is patented: Patent US 10578,620 B2: Methods for detecting molecules in samples (Patent Rights held by The Francis Crick Institute).

**Slide Preparation and COMET seqIF:**

FFPE slides were de-waxed and antigen retrieval was performed by placing the slides in Dewax and HIER Buffer H (Epredia, catalogue number: TA-999-DHBH) inside the PT Module (Epredia, catalogue number: A80400011) for 20 mins at 95°C. Processed slides were immediately placed into Multi-staining Buffer (Lunaphore Technologies SA, catalogue number: BU06) until use. Slides were transferred to the COMET instrument where a microfluidic COMET Chip (Lunaphore Technologies SA, catalogue number: MK03) was sealed on top of the processed FFPE tissue section by the COMET instrument to form a closed reaction chamber. FFPE tissue sections were stained by reagents delivered via the microfluidic channels according to the pre-loaded seqIF protocols. Automated seqIF protocols were generated using the COMET Control software and consisted of iterative autofluorescence, staining, imaging and elution cycles[17].

Primary antibodies, secondary antibodies and nuclear counterstain Hoechst 33342 were diluted to their desired concentrations using multi-staining buffer (MSB, Lunaphore Technologies SA, catalogue number: BU06). Slides were imaged at 20X magnification within a 12.5 x 12.5mm imaging area by the integrated fluorescent microscope using DAPI (exposure time: 25 ms), TRITC (exposure time: 250 ms) and Cy5 (exposure time: 400 ms) channels for every imaging cycle. Primary and secondary antibodies were incubated for 8 and 16 minutes respectively in each cycle based on prior optimization. After each imaging cycle, antibodies were eluted using the default settings for each elution cycle. Once completed, the COMET seqIF protocol generated a multi-layer OME-TIFF consisting of the stitched and aligned images from each imaging cycle[17].

**Photophysical parameters for quantification of protein interactive states.**

As input the algorithm takes a lifetime image of donor in the presence of acceptor ($\tau_{DA}$) and a lifetime image of the donor ($\tau_D$), followed by calculation of reduction of $\tau_{DA}$ compared to $\tau_D$, which is reflected in a metric called FRET-efficiency:

$Ef = [1 - <\tau_{DA}>/<\tau_D>]x100$ **Eq-1**

FRET-efficiency (Eq-1) is calculated as an average for each coincident region. Eq-1 is directly related to the distance between the donor and acceptor fluorophores (Atto488 and Alexa 594) (Eq-2), where r is the distance between Atto488 and Alexa594 in these experiments. '$R_0$', the Förster radius in this case, is 5.83nm and it is the distance at which the transfer efficiency is 50%.

$Ef = \frac{R_0^6}{R_0^6+r^6}$ **Eq-2**

For each region of coincidence in the two-site assay, we computed the mean distance between the donor and the acceptor.

$r = \sqrt[6]{R_0^6 * \frac{(1-Ef)}{Ef}}$ **Eq-3**

Each coincident region is color-coded based on a colourmap that reflects a range of 0 to 50% FRET efficiency. This color scale is used to create a heatmap, which is mapped automatically on the donor expression or the COMET expression image, in this case reflecting PD-1 expression.

**Time-Resolved intercellular/immune FRET (iFRET) determined by frequency-domain FLIM.**

The quantitative molecular imaging platform utilizes a custom-made semi-automated frequency-domain FLIM (Lambert Instruments). The first slide (donor only) is excited by a modulated (40MHz) diode 473nm laser, and the lifetime of the donor alone recorded. The second slide was excited by the diode modulated 473nm laser and lifetime of the donor in the presence of the acceptor recorded. The reduction of donor lifetime (caused by resonance energy transfer) due to the presence of the acceptor reports on distances of 1-10nm and therefore acts as a “spectroscopic ruler” enabling to quantify receptor-ligand protein interactions. We identify the coincidence regions where both the donor and acceptor are observed and their Ef values were automatically calculated and mapped on to the expression levels of PD-1.
