## Supplementary Table 1 for "Functional Spatial Mapping of the Tumor Immune Microenvironment In Advanced Melanoma Patients"

**Table 1. Two-site Assays Used to Quantify Immune Checkpoint Interactions**

| Reference: | One or Two-site assay(s) used: | Assay Range: | Protein Interaction quantified: | Does assay directly quantify functional interaction: | How are interactions scored: |
|---|---|---|---|---|---|
| Naei, V.Y., Tubelleza, R., Monkman, J. et al. J Transl Med 23, 177 (2025). https://doi.org/10.1186/s12967-025-06186-y | Proximity Ligation Assay (PLA) | Within 40nm | PD-1/PD-L1 | No, proximity of PD-1+ and PD-L1+ cells within 40nm are inferred as interacting | Interactions classified as positive or negative based on PD-1/PD-L1 intensity thresholds |
| Roche VENTANA PD-L1 (SP142) assay | One-site Immunohistochemistry (IHC) for PD-L1 | 70-100nm | N/A | No, the proportion of tumor (TC) or immune (IC) cells are scored as the proportion of viable cells showing PD-L1 staining of any intensity | N/A |
| Veeriah S., Leboucher P., de Naurois J. et al. Cancer Res (2014) 74 (18): 4983–4995. https://doi.org/10.1158/0008-5472.CAN-13-3382 | Coincidence amplified intracellular Förster Resonance Energy Transfer (aFRET) | 1-10nm | Intracellular Activation of protein kinase B (Akt) by Phosphorylation | No, quantifies intracellular Akt activation status based on interaction between pan-Akt and phospho-Akt antibodies within 10nm | N/A |
| Sánchez-Magraner L., Miles J., Baker C. L. et al. Cancer Res (2020) 80 (19): 4244–4257. https://doi.org/10.1158/0008-5472.CAN-20-1117 | Intercellular/immune Förster Resonance Energy Transfer (iFRET) and PLA | 1-10nm, 17-40nm | PD-1/PD-L1, CTLA-4/CD80 | PLA: no, see above<br><br>iFRET: yes, PD-1/PD-L1 interactions based on chromophore energy transfer between cells within 10nm | Median FRET efficiency (Ef) across tissues |
| Sánchez-Magraner L., Gumuzio J., Miles J., et al. J Clin Oncol (2023) 41 (14): 2561-2570. https://doi.org/10.1200/JCO.22.01748 | QF-Pro (based on iFRET) | 1-10nm | PD-1/PD-L1 | Yes, see above | Median Ef values across tissues |
| Sarygina E, Applebee C., McIntyre A., et al. BJC Reports (2024) 2(1):1-11. https://doi.org/10.1038/s44276-023-00033-7 | Functional Oncology Mapping (FuncOmap) | 1-10nm | PD-1/PD-L1 | Yes, iFRET coupled to a computational algorithm for automated Ef thresholding. | Per-pixel Ef values per coincidence regions across patient tissue samples |
