## Supplementary Table 2 for "Functional Spatial Mapping of the Tumor Immune Microenvironment In Advanced Melanoma Patients"

| Patient # | Gender | Age (Years) | Ethnicity | Prior Therapy | Neoadjuvant ICI Regimen Received | Cycles Given | TNM Stage | AJCC Stage | Pathological Response | RVT (%) | Tumor Necrosis (%) | Tumor Melanosis/Fibrosis (%) | PFS | OS | Status |
|---|---|---|---|---|---|---|---|---|---|---|---|---|---|---|---|
| 15 | F | 69 | Hispanic | No | Anti-PD-1 + Anti-LAG3 | 2 | pT3b, pN2b, cM0 | IIIC | NR | 70 | 25 | 25 | 63 | 63 | Alive |
| 27 | M | 59 | Non-Hispanic | No | Anti-PD-1 + Anti-LAG3 | 2 | pT1a, cN2b, cM0 | IIIB | NR | 60 | 35 | 5 | 62 | 62 | Alive |
| 34 | F | 74 | Non-Hispanic | No | Anti-PD-1+ Anti-CTLA4 | 5 | pT4b, pN2a, pMx | IIIC | PR | 50 | N/A | N/A | 26 | 36 | Deceased |
| 38 | M | 71 | Non-Hispanic | No | Anti-PD-1 + Anti-LAG3 | 2 | cN2b, cM0 | IIIB | Near-CR | 3 | 10 | 87 | 61 | 61 | Alive |
| 49 | M | 51 | Non-Hispanic | No | Anti-PD-1 + Anti-LAG3 | 1 | pT4b, pN1c, pMx | IIIC | Near-CR | 5 | <5 | 92 | 50 | 50 | Alive |
| 53 | F | 57 | Non-Hispanic | No | Anti-PD-1 + Anti-LAG3 | 2 | pT1b, cn2b, cM0 | IIIB | CR | 0 | 70 | 30 | 45 | 45 | Alive |
| 65 | M | 71 | Non-Hispanic | No | Anti-PD-1 + Anti-LAG3 | 3 | rpT0, pN1b, cM0 | IIIB | CR | 0 | <5 | 95 | 46 | 46 | Alive |

**Supplementary Table 2. Clinicopathologic Features of Patients:** TNM and AJCC stages were categorized according to the American Joint Committee on Cancer (AJCC) ($8^{th}$ Edition). Pathologic Responses were categorized according to International Neoadjuvant Melanoma Consortium (KNMC) guidelines. Abbreviations: RVT; Residual Viable Tumor, PFS; Progression Free Survival, OS; Overall Survival.
